# The local anaesthetic procaine prodrugs ProcCluster^®^ and Procaine-hydrochloride impair SARS-CoV-2 replication *in vitro*

**DOI:** 10.1101/2021.06.07.447335

**Authors:** Clio Häring, Josefine Schroeder, Bettina Löffler, Beatrice Engert, Christina Ehrhardt

**Affiliations:** Section of Experimental Virology, Institute of Medical Microbiology, Center for Molecular Biomedicine (CMB), Jena University Hospital, Jena, Germany; Institute of Medical Microbiology, Jena University Hospital, Jena, Germany; inflamed pharma GmbH, Jena, Germany

**Author notes:** corresponding author Christina Ehrhardt, Section of Experimental Virology, Institute of Medical Microbiology, Hans-Knoell-Str. 2, 07745 Jena.

**Keywords:** SARS-CoV-2, influenza A virus, ProcCluster^®^, Procaine-hydrochloride, drug repurposing

## Abstract

The SARS-CoV-2 pandemic has had the world in suspense for more than a year. Even if more and more vaccines are approved there is still an urgent need for efficient antiviral treatment strategies. Here, we present data on the inhibitory effect of the local anaesthetic procaine, especially the prodrugs ProcCluster^®^ and Procaine-hydrochloride on SARS-CoV-2 infection *in vitro*. Remarkably, similar effects could be shown on the replication of influenza A viruses in cell culture systems. Since the active ingredient procaine is well-tolerated and already used in the clinics for anaesthetic purposes, the further investigation of this substance could enable its reuse in antiviral therapy, including SARS-CoV-2.

## Introduction

The ongoing spread of the novel severe acute respiratory syndrome coronavirus 2 (SARS-CoV-2), causing the coronavirus disease 2019 (COVID-19), is characterized by an enormous increase in infected people worldwide [1]. Comparable to previous pandemics caused by influenza A viruses (IAV), SARS-CoV-2 primarily targets the bronchial and alveolar epithelial cells. So far, treating severe cases of the common flu and COVID-19 is an unsolved challenge. Hallmarks of severe COVID-19 are pneumonia, pulmonary edema, acute respiratory distress syndrome (ARDS) and multiple organ failure, probably due to viremia [2]. Many patients showed cardiovascular complications accompanied by vascular abnormalities, high inflammatory markers and markers of cell-disturbance [3]. These data point to massive problems in the regulation of innate immune responses. Recently vaccine production and administration have increased and vaccination is the most effective method of protection. However, vaccinations would not confer protection against newly emerging virus subtypes and some persons cannot or will not be vaccinated. Accordingly, the identification of effective pharmaceutic ingredients is needed for infected persons and the repurposing of approved substances that are in use against other virus infections, or diseases, seems to be the appropriate method [4]. Besides targeting the virus itself, novel therapeutic strategies include the inhibition of virus-supportive cellular factors, or excessive immune responses, which result in tissue damage and inflammation.

Here, we investigated if procaine, which is usually used as a local anaesthetic, represents a potential candidate for further antiviral studies. Local anaesthetics are described to exhibit anti-inflammatory and antioxidative properties, among others [5]. A few studies further indicate antiviral potential of the substance against different viruses [6,7], others report regulatory functions on cellular factors, such as G protein-coupled receptors (GPR) [8], and the mitogen-activated protein kinases (MAPK) [9]. Notably, GPRs and MAPKs mediate virus-induced antiviral cellular functions, but are also used by viruses for own purposes to ensure efficient replication [10,11].

ProcCluster^®^ (PC) is a patented substance (PCT/EP2018/074089; EP21157974.3), also known as Procainum-hydrogen-carbonate (ProcHHCO_3_ * NaCl), which was manufactured and supplied by inflamed pharma GmbH (Jena). PC is based on procaine, an ester-type local anaesthetic, which is stabilized by the mineral salt enhancing the bioavailability. Since 2008 PC has been used in Germany for the preparation of prescription drugs, which are administered orally and dermally. Since 2012, the license to manufacture parenteral drugs as part of the permit-free inhouse production exists (§13 2b AMG). Similarly, Procaine-hydrochloride (ProcHCl) is a prodrug of procaine and is administered parenterally.

Thus, we analyzed if procaine, in particular PC and ProcHCl exhibit antiviral effects on SARS-CoV-2 and influenza A virus (IAV) infection *in vitro*.

## Results and Discussion

### ProcCluster^®^ and Procaine-hydrochloride reduce the replication efficacy of SARS-CoV-2 and influenza A virus in vitro

The potential antiviral effect of procaine on SARS-CoV-2 (Figure 1) and IAV (Supplemental Figure 1 (S1)) was examined by analysis of the virus-induced cell disturbance, viral mRNA synthesis, viral protein expression and progeny infectious virus particles. Detailed information on material and methods is provided in the Supplemental Figure 2 (S2) and previous studies [12,13].

**Figure 1.**
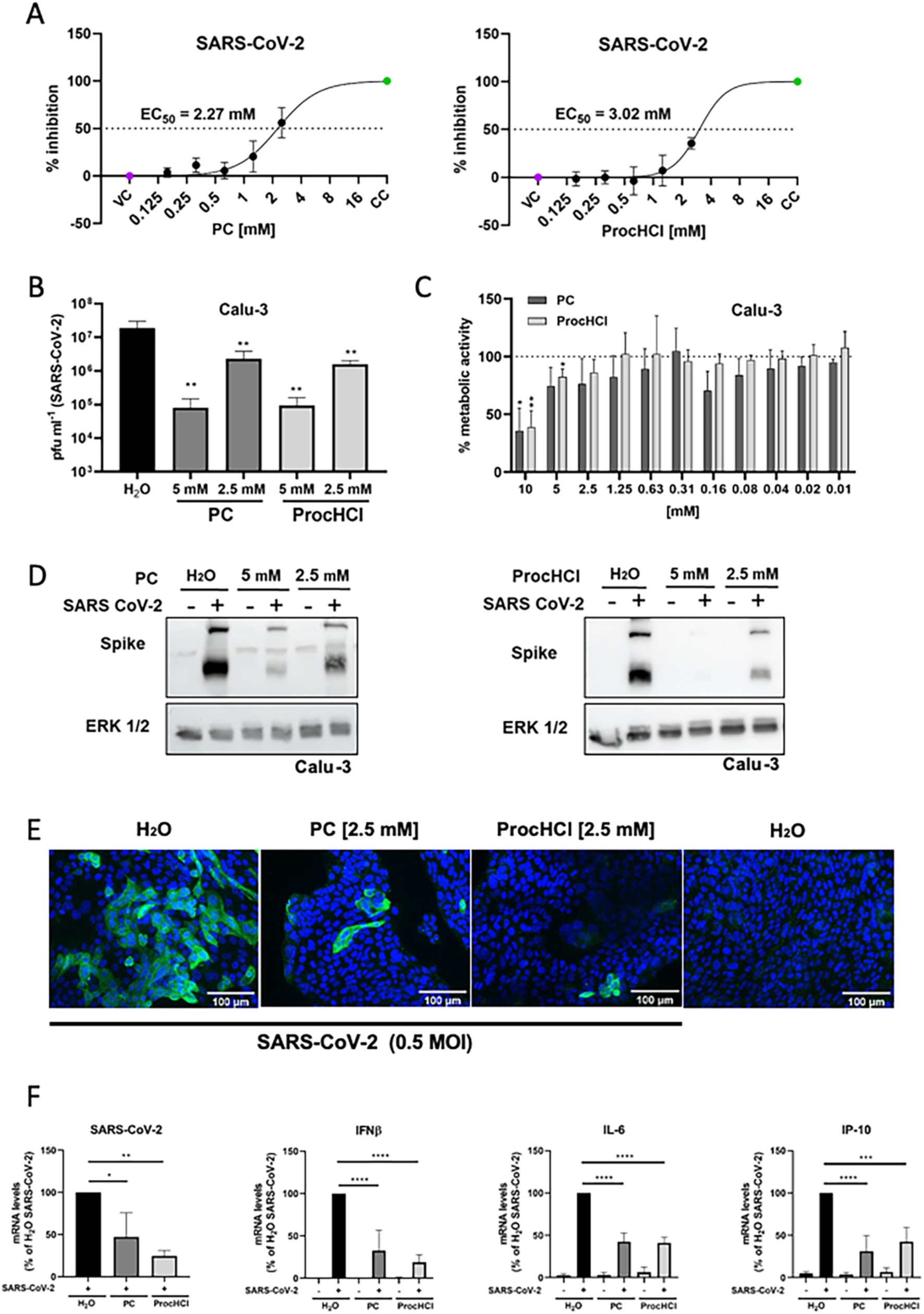
ProcCluster^®^ and Procaine-hydrochloride inhibit SARS-CoV-2 infection *in vitro*. Vero-76 cells (A) or Calu-3 cells (B-F) were infected with SARS-CoV-2 (A-B, D-F) or left uninfected (C) in absence and presence of the indicated substances (A-F). (A) The relative amount of surviving cells was measured to determine the effective concentrations 50 % (EC_50_) of PC and ProcHCl. (B) Virus titres were examined by standard plaque assays. (C) Proliferation of Calu-3 cells was analysed by MTT assay. (D) Protein synthesis of SARS-CoV-2 spike protein was visualized by western-blotting and equal protein load was verified by detection of the housekeeping protein ERK 1/2. (E) Immunofluorescence microscopy shows SARS-CoV-2 spike protein expression (green) and the nuclei were stained with Hoechst-33342 (blue). (F) The mRNA synthesis of SARS-CoV-2 (N1), cellular IFN-β, IL-6 and IP-10 were quantified by qRT-PCR. Data represent the mean + SD of three (A, B, F) or four (C) independent experiments, including two technical samples (*p < 0.05; **p < 0.01; *** p < 0.001; **** p < 0.0001).

Initially, we determined the effective concentration 50% (EC_50_) of the substances in Vero-76 or MDCK cells infected with the SARS-CoV-2 isolate SARS-CoV-2/hu/Germany/Jena-vi005159/2020 (SARS-CoV-2) (1 multiplicity of infection (MOI)) (Figure 1A) or influenza virus A/Puerto Rico/8/34 (IAV) (0.01 MOI) (Figure S1A) at 48 h post infection (p.i) by crystal violet staining. Here, procaine inhibited virus-induced cell death at millimolar concentrations [SARS-CoV-2: EC_50_ value of 2.27 mM (PC) and 3.02 mM (ProcHCl); IAV: EC_50_ value of 3.36 mM (PC) and 2.85 mM (ProcHCl)], implying reduced viral replication and spread. Afterwards, we examined the influence of procaine on progeny virus titres of SARS-CoV-2-infected Calu-3 cells (0.5 MOI) and IAV-infected A549 cells (0.1 MOI) in multi-replication-cycle experiments at 24 h p.i. (Figure 1B, D-F; S1B, D-F). Both, SARS-CoV-2 and IAV progeny virus titres were reduced in a concentration-dependent manner (Figure 1B, S1B). To exclude that the described effect on viral propagation is due to impaired cell viability, MTT-based proliferation assays were performed in the respective cell-lines (Figure 1C, S1C). As reduction of the substrate MTT only occurs in living and metabolically active cells, cellular proliferation and viability was determined. The results indicate that neither PC nor ProcHCl significantly affect cellular proliferation and viability. Thus, the observed impact of the substances on SARS-CoV-2 and IAV infection occurs independent of effects on cell viability. In accordance with the reduced progeny virus titres procaine-treatment of infected cells resulted in reduced SARS-CoV-2 spike protein levels in a concentration-dependent manner, as detected by western-blot analysis (Figure 1D). Correlating, a decreased production of the viral matrix 1 protein (M1) could be observed in IAV-infected A549 cells (Figure S1D). In order to visualize the extent of SARS-CoV-2 infection in Calu-3 cells and IAV infection in A549 cells in absence or presence of procaine, immunofluorescence microscopy studies were performed at 24 h p.i. (Figure 1E, S1E). In untreated, SARS-CoV-2-infected samples accumulation of spike protein is visible, which is reduced in procaine-treated samples. Accordingly, accumulation of the IAV nucleoprotein (NP) is visible in the cytoplasm and the nucleus of infected cells indicating ongoing replication, which is reduced in presence of procaine. These results point to a procaine-mediated inhibition of viral infection. Since one hallmark of severe disease of COVID-19 but also flu is enhanced cytokine expression accompanied with detrimental inflammation and cell death, the impact of both substances on viral mRNA synthesis as well as SARS-CoV-2- and IAV-mediated cytokine and chemokine expression were determined by qRT-PCR analysis at 24 h p.i,. As expected the treatment with procaine resulted in reduced viral mRNA synthesis of SARS-CoV-2 (N1) and IAV (M). In line, mRNA synthesis of different cytokines (IFNβ, IL-6, IP-10) (Figure 1F) or (MxA) (Figure S1F) was reduced in presence of procaine in comparison to untreated, infected cells.

In summary, our data show that PC as well as ProcHCl have antiviral activity against SARS-CoV-2 and IAV infection and also affect virus-induced cytokine expression. As mentioned above the procaine has a variety of properties. Future studies need to elucidate the procaine-, and in particular the PC- and ProcHCl-mediated effects on virus-induced cellular mechanisms.

## Acknowledgments

We thank Stefanie Kynast, Jessica Lux and Heike Urban for excellent technical support.

## Author contributions

CH, BL, BE and CE conceived and designed research. CH and JS performed experiments. CH, JS and CE analysed data. CH, JS and CE wrote the manuscript. All authors read and approved the manuscript.

