## supplemental material for "The local anaesthetic procaine prodrugs ProcCluster^®^ and Procaine-hydrochloride impair SARS-CoV-2 replication *in vitro*"

Supplementary Figure 1: ProcCluster® and Procaine-hydrochloride inhibit IAV infection *in vitro*

Supplementary Figure 2: Schematic representation of the infection and treatment procedure.

### Supplementary Figure 1

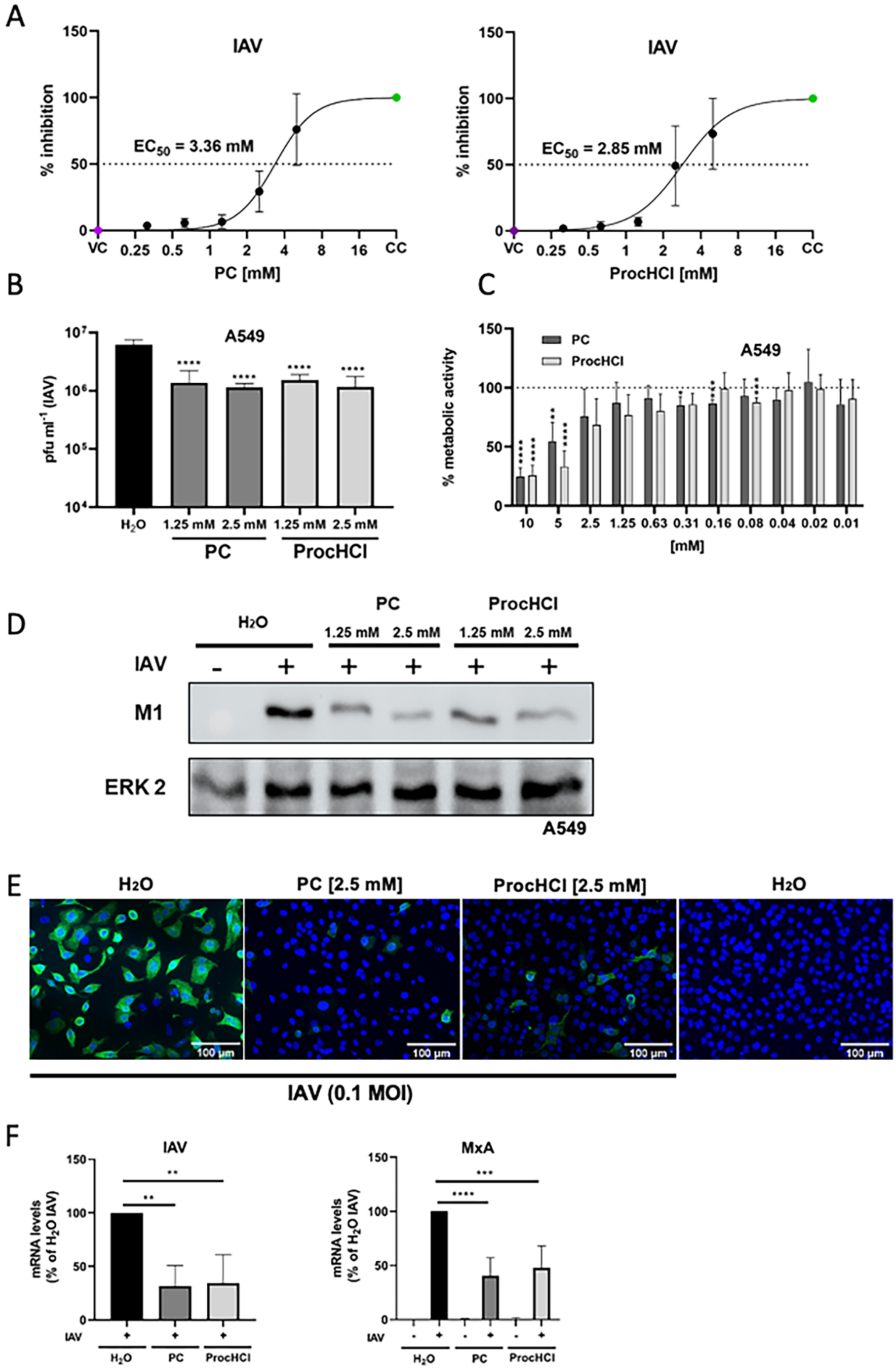

**Supplementary Figure 1 (S1):** ProcCluster® and Procaine-hydrochloride inhibit IAV infection *in vitro*.

MDCK cells (A) or A549 cells (B-F) were infected with IAV (A-B, D-F) or left uninfected (C) in absence and presence of the indicated substances (A-F). (A) The relative amount of surviving cells was measured to determine the effective concentrations 50 % ($EC_{50}$) of PC and ProcHCl. (B) Virus titres were examined by standard plaque assays. (C) Proliferation of A549 cells was analysed by MTT assay. (D) Protein synthesis of IAV matrix protein (M1) was visualized by western-blotting and equal protein load was verified by detection of the housekeeping protein ERK 2. (E) Immunofluorescence microscopy shows IAV nucleoprotein (NP) expression (green) and the nuclei were stained with Hoechst-33342 (blue). (F) The mRNA synthesis of IAV (M), and cellular MxA were quantified by qRT-PCR. Data represent the mean + SD of four (A, B, F) or five (C) independent experiments, including two technical samples (**$p < 0.01$; *** $p < 0.001$; **** $p < 0.0001$).

.

### Supplementary Figure 2

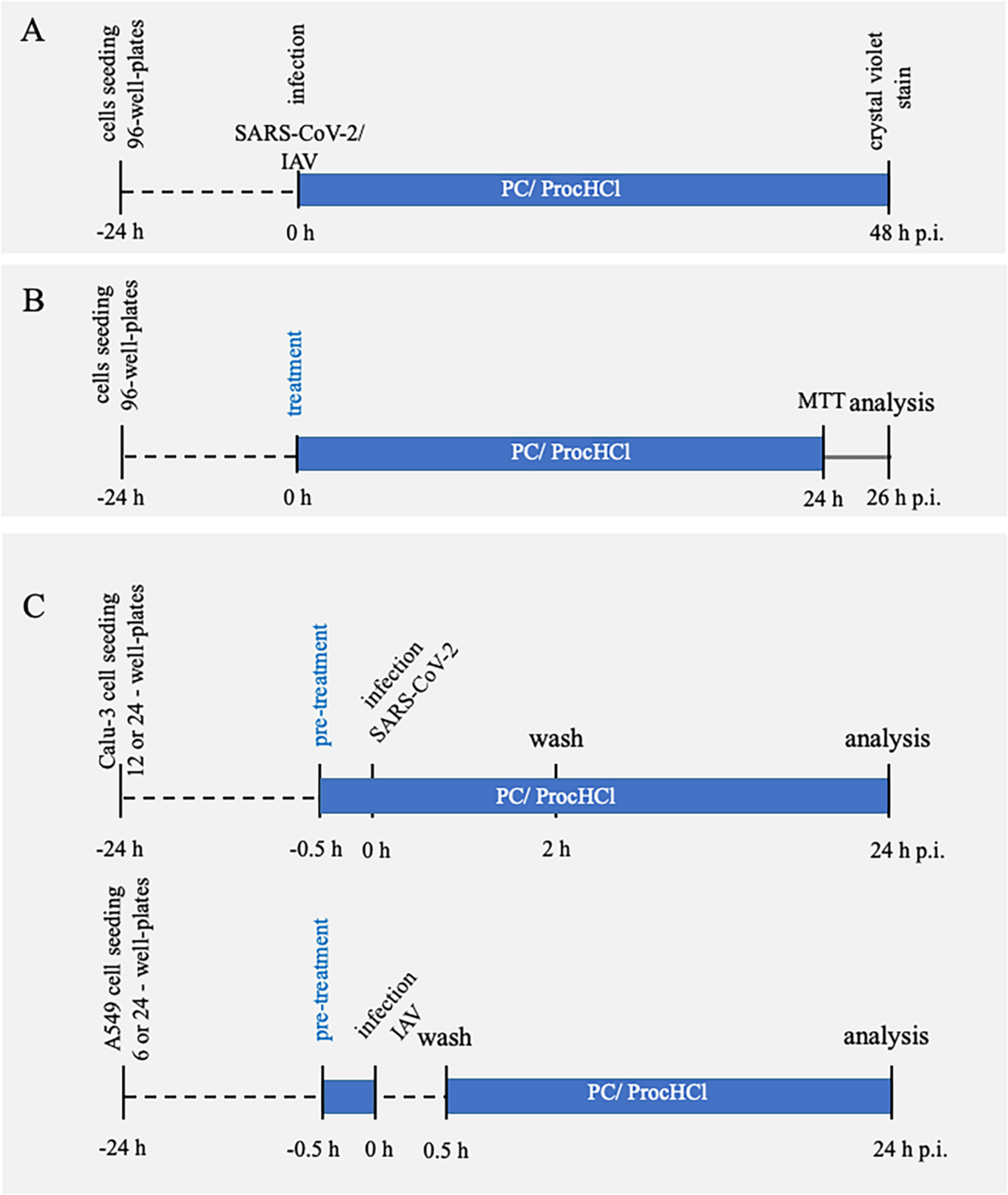


**Supplementary Figure 2 (S2):** Schematic representation of the infection and treatment procedure.

Vero-76, Calu-3- and A549 cells were cultivated in DMEM (10% FCS), and MDCK cells were cultivated in EMEM (10% FCS) at 37°C and 5% $CO_2$.

**Effective concentration 50% ($EC_{50}$)** (A) Vero-76 cells (for SARS-CoV-2: SARS-CoV-2/hu/Germany/Jena-vi005159/2020) or MDCK cells (for IAV: A/Puerto Rico/8/34) were seeded into 96-well plates the day prior to the experiment. Cells were washed and infected with SARS-CoV-2 (1 MOI) in DMEM (10% FCS) or IAV (0.01 MOI) in EMEM/BA (0.2 % BSA, 1 mM $MgCl_2$ and 0.9 mM $CaCl_2$; 100 U/ml Pen/Strep, 167 ng $ml^{-1}$ Trypsin-TPCK) and simultaneously treated with ProcCluster® (PC) (inflamed pharma GmbH, Jena, Germany) or Procaine-hydrochloride (ProcHCl) (Caesar & Loretz GmbH, Hilden, Germany) at the indicated concentrations. The cells were incubated at 37°C and 5% $CO_2$ for 48 h, after which the medium was removed, the cells were washed with PBS and fixed with crystal violet solution (0.2% crystal violet, 20% EtOH, 3% formaldehyde) over night (for SARS-CoV-2) or 20 min (for IAV). The cells were then washed with PBS several times and supplemented with DMSO. The $OD_{595}$ was measured and the relative absorbance of infected treated cells in comparison to uninfected untreated cells (100%) as well as infected untreated cells (0%) was used to determine $EC_{50}$ values by non-linear fit (least squares; variable slope)(GraphPadPrism 8.3).

**Cell Proliferation Assay (MTT)** (B) Calu-3 cells or A549 cells were seeded into 96-well plates 24 h prior to the experiment. The cells were left untreated or treated with a serial 1:2 dilution of PC or ProcHCl for 24 h, after which 25 µl MTT (Sigma Aldrich) was added and cells were incubated a further 2 h. The supernatant was carefully removed and DMSO was added. Metabolic activity was determined by relative absorbance at $OD_{562}$ in comparison to the solvent ($H_2O$)-treated control. Statistical significance was determined using multiple t-tests with Holm-Sidak method for correction.

**Viral infection and investigation via standard plaque assay, western-blot analysis, immunofluorescence studies and qRT-PCR:** (C) Calu-3 cells (for SARS-CoV-2) and A549 cells (for IAV) were pre-treated with PC, ProcHCl or solvent $H_2O$ 30 min prior to infection.

Cells were washed and infected in DMEM (10% FCS) (for SARS-CoV-2: 0.5 MOI) or PBS/BA (for IAV: 0.1 MOI) containing PC, ProcHCl or solvent control. After a second washing step at 30 min (IAV) or 2 h (SARS-CoV-2) the cells were incubated in fresh medium containing PC, ProcHCl or solvent control for 24 h p.i.. For IAV infection of A549 cells 167 ng $ml^{-1}$ Trypsin-TPCK was added to the medium. Supernatant was used to determine progeny virus titres by standard plaque assay.

**For plaque assays** Vero-76 cells (for SARS-CoV-2) or MDCK cells (for IAV) were seeded into 6-well plates to create a confluent cell layer. The cells were infected with serial dilutions of a sample in PBS/BA (0.2 % BSA, 1 mM $MgCl_2$ and 0.9 mM $CaCl_2$) supplemented with 100 U/ml Pen/Strep for 90 min (SARS-CoV-2) or 30 min (IAV). The inoculum was then replaced with EMEM supplemented with 0.2 % BSA, 0.01 % DEAE Dextran (Pharmacia Biotech, Germany), 0.2% NaHCO3 (Biozym), 100 U/ml Pen/Strep and 0.9 % agar (Oxoid, Wesel, Germany) and incubated at 37°C and 5 % $CO_2$ for 3 days. For IAV infection of MDCK cells 250 ng $ml^{-1}$ Trypsin-TPCK was added to the medium. Plaques were visualized by neutral red staining. Statistical significance was determined by ordinary one-way ANOVA with Dunnett's multiple comparisons test.

**For western-blot analysis** cells were lysed with Triton lysis buffer (TLB; 20 mM Tris-HCl, pH 7.4; 137 mM NaCl; 10% Glycerol; 1% Triton X-100; 2 mM EDTA; 50 mM sodium glycerophosphate, 20 mM sodium pyrophosphate; 5 µg $ml^{-1}$ aprotinin; 5 µg $ml^{-1}$ leupeptin; 1 mM sodium vanadate and 5 mM benzamidine) for 1 h. Cell lysates were centrifuged, and 5x Lämmli buffer (10 % SDS, 50 % glycerol, 25 % 2-mercaptoethanol, 0.02 % bromophenol blue, 312 mM Tris 6.8 pH) was added (1:5 dilution) and the mixture was boiled at 95°C for 10 min. Samples were subjected to SDS-PAGE and blotting. SARS-CoV-2 spike protein was detected using a rabbit polyclonal anti-SARS-CoV-2 spike S2 antibody (Sino Biological). Equal protein load was verified using a rabbit polyclonal anti-ERK1/2 (Cell Signaling) antibody. IAV M1 protein was detected using a monoclonal antibody (BioRad) and equal protein load was confirmed using a monoclonal anti-ERK 2 (Santa Cruz Biotechnology) antibody.

**For qRT-PCR** RNA from Calu-3 and A549 cells was isolated using the RNeasy Mini Kit (Qiagen, Hilden, Germany) according to the manufacturer's instruction. The QuantiNova Reverse Transcription Kit (Qiagen, Hilden, Germany) was used for cDNA synthesis from 400 ng of total RNA. For qRT-PCR the QuantiNova SYBR Green PCR Kit (Qiagen, Hilden, Germany) was used. Cycle conditions were set as follows: 95°C for 2 min, followed by 40 cycles of 95°C for 5 sec and 60°C for 10 sec. The qPCR cycle was ended by a stepwise temperature-increase from 60°C to 95 °C (1°C every 5 sec). The following primers were used: 2019-nCoV_N1_fw: 5'-GACCCCAAAATCAGCGAAAT-3'; 2019-nCoV_N1_rev: 5'-TCTGGTTACTGCCAGTTGAATCTG-3'; IAV_fw (Seg 7): 5'-CAATACGACCAAATCCGTTGAC-3'; IAV_rev (Seg 7): 5'-AGGGCATTTTGGACAAAGCGTCTA-3'; human_GAPDH_fwd: 5'-CTCTGCTCCTCCTGTTCGAC-3'; human_GAPDH_rev: 5'-CAATACGACCAAATCCGTTGAC-3'; human_IFNβ_fwd: 5'-ATGACCAACAAGTGTCTCCTCC-3'; human_IFNβ_rev: 5'-GGAATCCAAGCAAGTTGTAGCTC-3' human_IL-6_fwd: 5'-CAGCCCTGAGAAAGGAGACATG-3'; human_IL-6_rev: 5'-GCATCCATCTTTTTCAGCCATC-3'; human_MxA_fwd: 5'-GAAGGGCAACTCCTGACAG-3'; human_MxA_rev: 5‘-GTTTCCGAAGTGGACATCGCA-3‘; human_IP-10_fwd: 5′-CCAGAATCGAAGGCCATCAA-3′; human_IP-10_rev: 5′-TTTCCTTGCTAACTGCTTTCAG-3′

Statistical significance was determined by ordinary one-way ANOVA with Dunnett's multiple comparisons test.

**For immunofluorescence studies** Calu-3 cells (for SARS-CoV-2) and A549 cells (for IAV) were seeded in 24-well plates on cover slips 24 h prior to infection. After infection, which was performed as described above cells were washed with PBS twice and then fixed with 4% formaldehyde solution for 30 min at 37°C for SARS-CoV-2-infected samples or 15 min at room temperature for IAV-infected samples. The samples were then washed with PBS three times to remove the formaldehyde. Afterwards, cells were permeabilized for 15 minutes using 250 µl of 0.1% Triton X-100 (in PBS) (Roth, Germany) and unspecific binding sites were blocked for 1 h using PBS/BSA (3%). Samples were then incubated with anti-SARS2-spike antibody (mouse) (GeneTex) or anti-IAV nucleoprotein (NP) antibody (mouse) (BioRad) diluted 1:100 in PBS/BSA (3%) for 2 h, washed again with PBS and afterwards incubated with Alexa Fluor 488 goat anti-mouse IgG polyclonal antibody (Dianova) diluted 1:200 in PBS/BSA (3%) was added for 1 h. The nuclei were stained with bisBenzimide H 33342 trihydrochloride (Hoechst-33342; Merck) diluted 1:1000 in PBS/BSA (3%). The samples were mounted on slides using fluorescence mounting medium (Agilent, Santa Clara, CA, USA). Image acquisition and editing was done by using an Axio Observer.Z1 microscope (Zeiss, Jena, Germany) and the softwae Fiji V 1.52b (ImageJ).
